# Identification of Antigen-Specific T Cell Receptors with Combinatorial Peptide Pooling

**DOI:** 10.1101/2023.11.28.569052

**Authors:** Vasilisa A. Kovaleva, David J. Pattinson, Guanchen He, Carl Barton, Sarah R. Chapin, Anastasia A. Minervina, Qin Huang, Paul G. Thomas, Mikhail V. Pogorelyy, Hannah V. Meyer

## Abstract

T cell receptor (TCR) repertoire diversity enables the antigen-specific immune responses against the vast space of possible pathogens. Identifying TCR-antigen binding pairs from the large TCR repertoire and antigen space is crucial for biomedical research. Here, we introduce *copepodTCR*, an open-access tool to design and interpret high-throughput experimental TCR specificity assays. *copepodTCR* implements a combinatorial peptide pooling scheme for efficient experimental testing of T cell responses against large overlapping peptide libraries, that can be used to identify the specificity of (or “deorphanize”) TCRs. The scheme detects experimental errors and, coupled with a hierarchical Bayesian model for unbiased interpretation, identifies the response-eliciting peptide sequence for a TCR of interest out of hundreds of peptides tested using a simple experimental set-up. Using *in silico* simulations, we demonstrate the varied experimental settings in which *copepodTCR* yields efficient and interpretable TCR specificity results. We validated our approach on a library of 253 overlapping peptides covering the SARS-CoV-2 spike protein, split across 12 pools. A single stimulation with combinatorial pools identified the correct epitope of two TCRs with known specificity and then deorphanized two SARS-CoV-2 associated TCRs shared among a large cohort of COVID-19 patients. We provide experimental guides to efficiently design larger screens covering thousands of peptides which will be crucial to identify antigen-specific T cells and their targets from limited clinical material.

## Introduction

Identifying antigen-specific T cell responses and their targets is crucial in numerous applications, including vaccine research^1^, cancer immunotherapy development^2^, and autoimmunity treatment^3^. A T cell’s ability to mount a targeted immune response is defined by the specificity of its T cell receptor (TCR). TCRs are generated semi-stochastically by somatic rearrangement of germline-encoded gene segments^4^. This process generates TCRs which can respond with high specificity to their cognate epitope. For conventional T cells, the cognate epitopes are short peptide sequences presented on major histocompatibility complexes (MHC). MHC molecules are present on the surface of all nucleated cells (MHC class I) and professional antigen-presenting cells (MHC class II).

While there are large efforts to predict peptide-MHC binding^5^ and TCR-epitope specificity computationally^6^, these are still limited by lack of unbiased, true positive and negative training data^7,8^ and also vary greatly depending on peptide-MHC context^9^. Thus, there is still a critical role for efficient, empirical T cell epitope discovery.

Identifying the peptide specificity of a given TCR requires an experimental assay in which T cells expressing the TCR of interest interact with antigen presenting cells that express candidate peptide-MHC complexes on their surface, coupled with a quantitative read-out of T cell activation upon this peptide-MHC encounter. An ideal experiment would allow high-throughput screening of a large peptide library against a given TCR, with mechanisms to detect experimental errors (false positives and false negatives), in a simple experimental setup. Careful peptide library design and unbiased interpretation are crucial to achieve this.

A peptide library is a sample of peptides of a particular length, that can be used in a TCR assay. To detect epitopes that activate the TCR, and provide robust negative results, the library should tile the whole protein or proteome space of potential antigenic peptides. Comprehensive coverage can be achieved by a ‘sliding window’ approach where the protein space of interest is sliced into overlapping peptides of defined length and overlap, that cover the whole space (Figure 1A). However, while optimal for coverage, the use of overlapping peptides introduces a significant challenge in the design and interpretion of a high-throughput T cell activation assay, particularly because recognized epitopes will be present in multiple peptides in the library.

**Figure 1.**
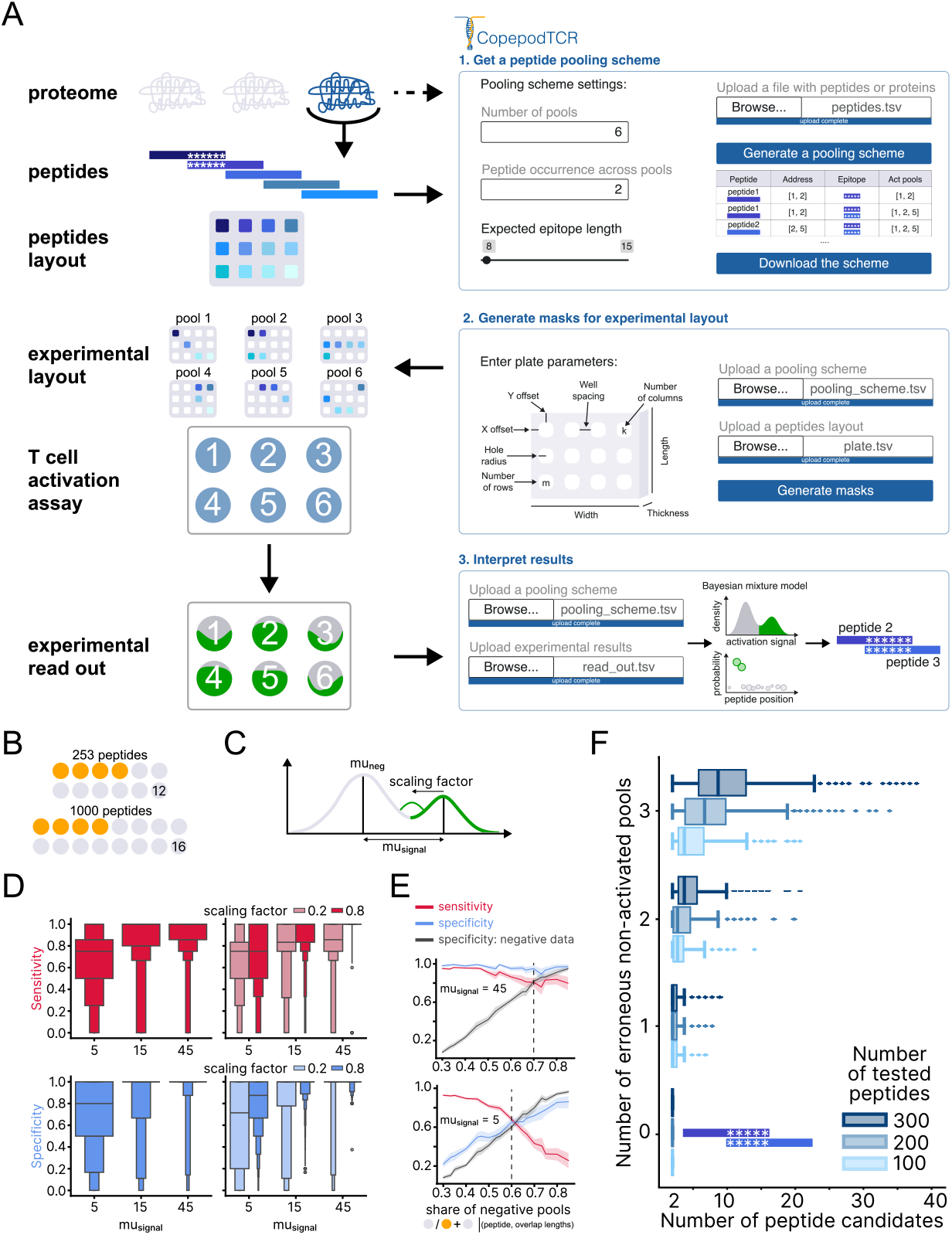
Combinatorial peptide pooling design. A. Experimental setup (left panels): synthetic peptides, with predefined overlap from a protein/proteome of interest, are mixed according to the pooling algorithm and tested for their ability to elicit a T cell response in a T cell activation assay. The results of the activation assay indicate a pair of overlapping peptides, with the overlapping section (asterisks) identified as the cognate antigen. Right panels show a schematic of the *copepodTCR* user interface, as well as an example activation model fit and results output. B. Examples of the peptide pooling setup, with the indicated number of peptides and pools (number in the last circle), where yellow circles indicate the number of pools an individual peptide is added to. C. Key parameters in simulation scheme for generation of synthetic data. Negative pools were sampled from a normal distribution with *μ* = *mu*_neg_; positive pools were generated by sampling from a derivative of the negative distribution, obtained by adding a *signal* with *mu*_signal_. To simulate additional variance in the positive distribution, one pool from the positive data was scaled to induce a shift toward the negative distribution. D. and E. Activation model predictions on synthetic data. D. Sensitivity and specificity of the activation model improve with increasing *mu*_signal_ and decreasing scaling factor. Results shown are from 12,960 independent simulations. E. Sensitivity and specificity are a function of the expected share of negative pools and the distance between the mean of the negative and positive pool intensities (*mu*_signal_). The share of negative pools equals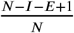, where *N* is the total number of pools, *I* is peptide occurrence (number of pools to which one peptide is added), and *E* is the number of peptides sharing one epitope. The vertical dashed line indicates the experimentally recommended share of negative pools. Negative data: specificity for simulations where all pools were sampled from the negative distribution. Results shown are from 12,636 independent simulations, including 4,212 negative data simulations. F. The number of possible candidates depends on the number of tested peptides and the number of erroneous non-activated pools. Results display a simulation with 18 pools and 6 pools per peptide and epitope presence in two consecutive peptides. For zero erroneously non-activated pools, the algorithm returns two peptides whose shared sequence contains the cognate epitope.

The simplest design, where each peptide is individually tested against each TCR, is time-consuming and reagent-intensive (e.g. as in^10^; Extended Data Fig 1A). An alternative approach, matrix pooling^11^, partially overcomes these limitations. In matrix pooling (Extended Data Fig 1B), a *n* × *m* experimental plate with a single peptide per well are combined into *n* row and *m* column peptide pools. In this design, each peptide is present in one row and one column, and a TCR-specific peptide should thus lead to the activation of exactly two pools. These can then be mapped back to identify the single activating peptide. Matrix pooling is more efficient compared to testing peptides individually as it requires less time and fewer reagents. Combinatorial peptide pooling (CPP) schemes using tables of addresses for combining peptides into pools^12,13^ (Extended Data Fig 1C) can further reduce the number of peptide pools required. In CPPs each peptide is added to a unique subset of pools (a “binary address”), which leads to specific pool activation patterns based on their combinations of peptides. The benefit of this approach is that a large number of peptides can be encoded by a low number of pools. However, for overlapping peptide libraries, a TCR will recognize more than one peptide which will make it hard to uniquely identify activating peptides.

Here, we developed *copepodTCR*– Combinatorial Peptide Pooling Design for TCR specificity –an algorithm for TCR deorphaning that addresses the three critical challenges in peptide library and experimental design for TCR specificity assays outlined above: it i) considerably reduces experimental resources and time required for testing by using a CPP approach, ii) accounts for overlapping peptides, and iii) enables the detection of errors in the experimental process and even in case of the error still significantly narrows down the list of peptide candidates. We demonstrate its ease of application and robustness of experimental design across a wide range of parameters, simulate erroneous experimental outcomes and their implications for experimental validation, and show its power in reducing experimental complexity. We experimentally validated our approach on a library of 253 overlapping peptides covering the SARS-CoV-2 spike organized into 12 pools, validating both known TCR targets and deorphanizing two TCRs of hitherto unknown specificity.

## Results

We have developed two independent computational models that jointly enable efficient design and interpretation of high-throughput TCR-pMHC specificity assays, which we describe below. Both models are implemented in the open-source Python package *copepodTCR*. For ease of use, we have developed a graphical user interface for the streamlined design and interpretation of custom experimental peptide pooling assays against any antigen.

### Efficient Design of high-throughput TCR-pMHC specificity assays

We previously developed a binary coding scheme that enables efficient construction of combinatorial pooling schemes for high-throughput experimental assays across biological applications^14^. These balanced constant-weight Gray codes for detecting consecutive positives (DCP-CWGCs) consist of a sequence of binary addresses, where a single address encodes the pooling scheme for a single item. Each position (bit) in the address indicates a pool, where 1 indicates the presence of, and 0 the absence of, the item in the pool. In the context of high-throughput TCR-pMHC specificity assays, items represent peptides, and pools are peptide mixtures added to antigen-presenting cells for peptide priming. This coding scheme is ideally suited to CPPs for high-throughput TCR-pMHC specificity assays, as it formalizes the requirements for peptide library compositions and makes their efficient design algorithmically tractable. We describe these requirements below and provide an intuition on the implementation. Details on the algorithm are given in the Supplementary Note.

### Unique addresses and uniqueness of their unions allow overlapping peptide design

Tiling the peptide space with overlapping peptides leads to epitope sharing in successive peptides, thus an epitope evoking T cell activation will do so in all pools successive peptides are present in. DCP-CWGCs provide a unique address for each peptide and can accommodate the detection of overlapping peptides by ensuring that the union of addresses for successive peptides is unique. This ensures that each peptide overlap results in a unique combination of activated pools, allowing for a clear interpretation of results.

### Error detection by maintaining a constant number of activated pools for any given epitope

In the T cell activation assay for a tiled peptide space, we expect that the T cell is activated by both of the two overlapping peptides containing the T cell’s cognate epitope. However, activation assays may yield false negative results if the activation in a given pool fails. False positives may occur due to pipetting errors, or if two epitopes in the tiled space both cause activation. DCP-CWGCs ensure that each peptide is added to the same number of pools. Any observed deviation in the number of activated pools therefore indicates a false positive or negative result.

### Balanced pool design for low and uniform error rates

T cell activation based on TCR-pMHC interaction is influenced by a number of factors including the TCR binding strength and the ‘hit rate’ of encountering the cognate antigen. While TCR binding strength will be specific for each TCR and cognate peptide, the hit rate can be influenced by ensuring sufficient peptide presentation to the tested T cells. Crucially, in multiplexed assays the hit rate should be constant across pools, to ensure consistent peptide exposure across experimental units. DCP-CWGCs ensure that the number of peptides per pool is consistent and balanced. Such balance not only results in uniform error rates across all pools but also minimizes the overall error probability, as the error rate is spread evenly among the pools.

### Unbiased interpretation of high-throughput TCR-pMHC specificity assays

We developed a hierarchical Bayesian model that can identify T cell-activating epitopes from the CPP assay based on a quantitative and continuous read-out of the T cell activation assay. Intuitively, we model the user-provided T cell activation read-out as the scaled sum of signal obtained from a negative and a positive distribution (Extended Data Fig 3). The model infers parameters of the negative distribution from negative controls, if they are included; otherwise, it infers them from the pool with the lowest activation signal. The distribution of the positive signal is modeled as an offset of the negative distribution, which can be interpreted as the addition of a signal to the underlying noise. The main output is the activation probabilities per peptide, visualized along the protein position. In addition, we return the list of the most probable cognate peptides and classification of pools into positive and negative distributions.

### Using *copepodTCR* to test TCR specificity against a large, tiling peptide library

The experimental setup (Figure 1A, left column) starts with defining the protein/proteome of interest and obtaining peptides tiling its space. This set of peptides, containing an overlap of a predefined length, is entered into the *copepodTCR* web application (Figure 1A, right column). *copepodTCR* then generates a peptide pooling scheme and, optionally, provides the pipetting scheme to generate the desired pools as either 384-well plate layouts or punch card models which could be 3D printed and overlaid on the physical plate or pipette tip box. Following this scheme, the peptides are mixed, and the resulting peptide pools are tested in a T cell activation assay. The activation of T cells is measured for each peptide pool with the assay of choice, such as flow cytometry- or microscopy-based activation assays detecting expression of a reporter gene. The experimental measurements for each pool are entered back into *copepodTCR* which employs a Bayesian mixture model to identify activated pools. Based on the activation patterns, it returns the set of overlapping peptides leading to T cell activation.

### *In silico* CPP assays inform experimental design and experimental error detection with *copepodTCR*

To guide the user in the parameter choice (Figure 1B) for designing CPP experiments with *copepodTCR*, we generated *in silico* experiments (Figure 1C) capturing a wide range of experimentally observable results including signal-to-noise ratio of read out (Figure 1C, *mu*_*signal*_) or strength of activation signal depending on the peptide context of the epitope (Figure 1C, scaling factor). For each of these settings, we varied the pooling design parameters (e.g. number of pools, peptide occurrence, number of tested peptides, peptide length) to guide experimental design and results interpretation. Full details on the generative model and the tested parameter ranges are in the methods.

Overall, the CPP design and model evaluation yielded consistent, highly specific and sensitive results across the wide range of user parameters that we assessed (see Extended Data Fig 5). As expected, the true positive (sensitivity) and true negative (specificity) rates of the model increased with the signal parameter, representing the signal-noise difference between negative and positive distributions (Figure 1D, first column). More consistent activation values across peptide contexts of the epitope, captured in a scaling factor, also resulted in better model performance (Figure 1D, second column), further confirming that the model’s performance improves when the difference in the signal between positive and negative pools is more pronounced.

We then investigated the impact of user design choices on the reliability of the model. Specifically, we evaluated the scenario of result interpretation as a function of the expected share of negative pools. This expected negative share is used as a prior in the model, representing the probability that a given pool comes from the negative distribution. It is calculated as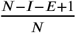, where *N* is the total number of pools, *I* is the peptide occurrence, and *E* is the number of peptides sharing one epitope. If the response-eliciting epitope is present in the peptide library and leads to T cell activation beyond background signal, specificity and sensitivity are consistently high across all tested negative shares (Figure 1E, mu_signal_ = 45). Importantly, for very weak activation signal, the negative share was a crucial user parameter choice for model performance (Figure 1E, mu_signal_ = 5), with decreasing sensitivity and increasing specificity for larger negative shares. Thus, for CPPs with higher uncertainty of the epitope presence in the peptide library, choosing a balanced negative share of 0.5-0.6 is recommended for best results. When a pronounced difference between positive and negative distributions is expected, i.e. there is a strong assumption that the target epitope is in the peptide library, an experimental setup with a higher negative share (approximately 0.7) and thus the overall cost can be reduced.

Next, we evaluated how the developed CPP scheme facilitates error detection. The peptide pooling scheme ensures that the number of activated pools remains constant for any given epitope (Figure 1F, number of erroneously non-activated pools equal to 0), thus any deviation of the observed number of activated pools from the expected can be detected as an error. When an error is detected, the tool effectively narrows down the list of potential peptide candidates responsible for the activation. The extent to which the list of candidates is reduced is influenced by the number of peptides tested and the number of erroneous non-activated pools. For instance, with the presence of one erroneous non-activated pool, the algorithm can refine the list of potential peptide candidates to approximately 10 (Figure 1F, number of erroneously non-activated pools equal to 1).

### *copepodTCR* CPP scheme successfully validates known TCR peptide specificity

To test that *copepodTCR*’s experimental design allows for the robust detection of true positive and negative TCR specificity, we designed CPP assays for two NFAT-GFP reporter Jurkat 76.7 TCR-transgenic cell lines with known peptide specificity. We first tiled and synthesized the SARS-CoV-2 spike protein into 253 17 amino acid (aa) long peptides with a 12 aa overlap between consecutive peptides. We mixed these peptides according to the CPP scheme generated by *copepodTCR* into 12 pools and performed T cell activation assays (in triplicates) for reporter lines stably expressing either TCR6.3 from ref.^15^ recognising the *S*_167−180_ epitope in context of DPB1*04:(01/02) or YLQ TCR 1 from ref.^16^ recognising the *S*_269−277_ epitope in context of HLA-A02+. We then measured the activation signal for each pool as the fraction of GFP+ (activated) T cells by both flow cytometry (Figure 2A) and fluorescence microscopy (Extended Data Fig 2). We used these activation signals as the input to the interpretation module of *copepodTCR* based on the Bayesian mixture model (Figure 2C), which enabled us to correctly identify the known, cognate peptides of the TCR6.3 and YLQ TCR 1 reporter cell lines (Figure 2D).

**Figure 2.**
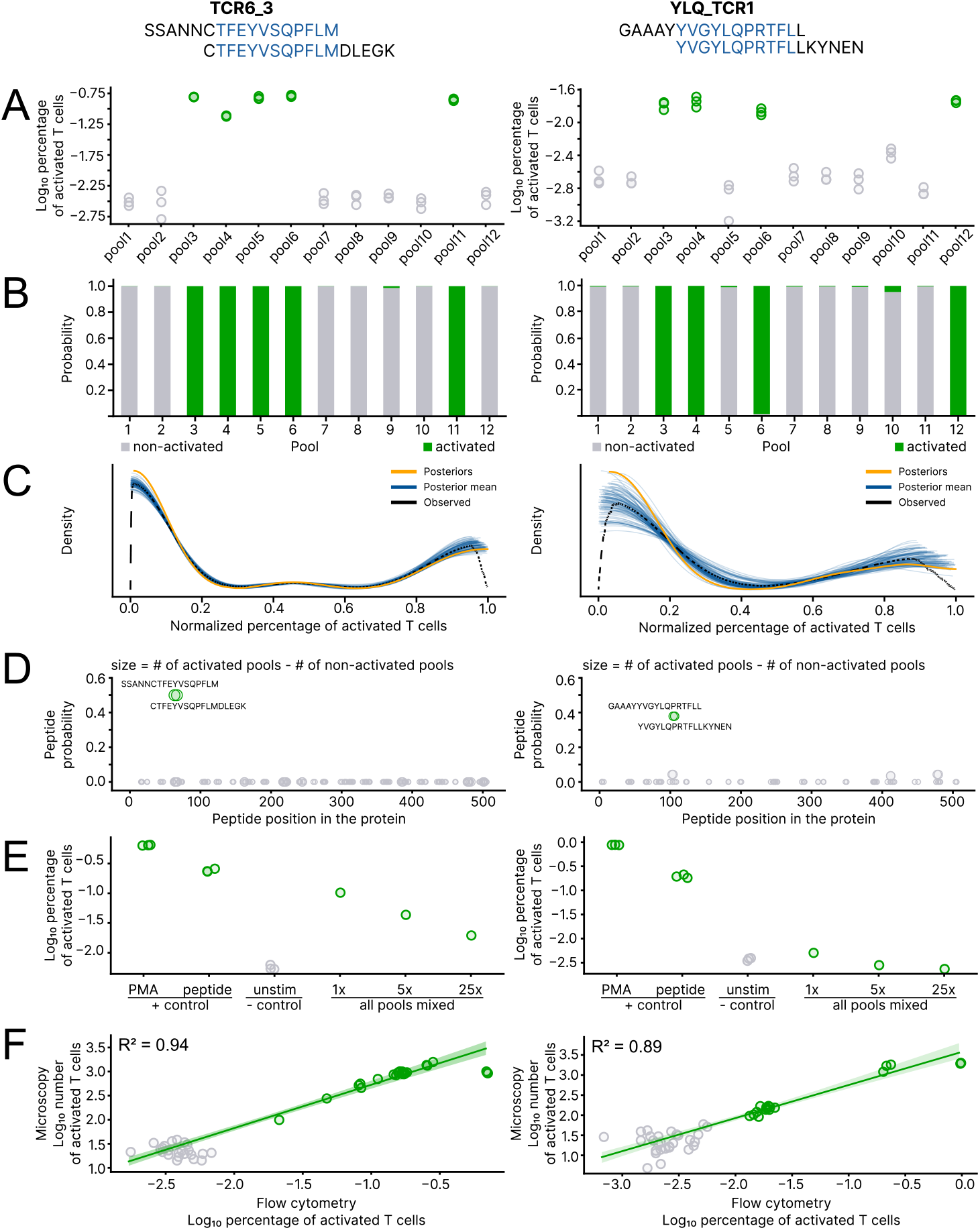
CopepodTCR design successfully recapitulates known TCR specificity. Experimental and model results of a CCP assay with *copepodTCR* for the T cell reporter cell lines expressing DPB1*04:(01/02)-restricted TCR6.3 (left column) and HLA-A02+ restricted YLQ-TCR1 (right column) with their known, cognate peptides indicated in blue letters in the peptide sequence. A. Flow cytometry read out as percentage of GFP-producing (activated) cells in the T cell activation assay (y-axis) for each pool in the design (x-axis), with pools detected as activated/non-activated in green and gray, respectively (determined from model fit in B.). B. Inferred probability (from C) of each measurement coming from the distribution of activated pools (green) and from a distribution of non-activated pools (gray). C. Bayesian mixture model (blue and orange) of observed (black) pool activation signal for the read out obtained by flow cytometry. D. Per-peptide probabilities calculated from B. The size of the circle shows the difference between the number of activated and non-activated pools to which a peptide was added. E. Sensitivity of the assay. Left: activation of the cell line by PMA/ionomycin, single cognate peptide and baseline activation levels without peptide stimulation. Right: The activation signal from all pools mixed together (1x, all pools mixed), and a 5-fold (5x), or 25-fold (25x) dilution thereof. All dilutions show activation levels above unstimulated negative control. F. Correlation of flow cytometry and microscopy-based read out of the T cell activation assay, each represented by percentage of activated cells and color-coded as in A.

To assess the sensitivity of the approach, we mixed all 253 peptides in a single pool and then diluted the mixture 5- and 25-fold. For the MHCII-restricted T cell line, the dilution caused a gradual decrease in the number of activated T cells, but the activation was still distinguishable when compared to the negative unstimulated control (Figure 2E, left panel). However, for the MHCI-restricted T cell line, the dilution resulted in an activation signal that was indistinguishable from the negative control (Figure 2E, right panel). These data suggest that pools with a much larger number of peptides and thus lower concentration of each individual peptide will still trigger detectable activation for MHCII-restricted T cell responses. Thus, it allows one to use *copepodTCR* for larger overlapping peptide libraries covering entire viral proteomes. Applying *copepodTCR* to the 1879 peptides tiling the SARS-CoV-2 proteome resulted in 18 pools, on average containing 625 peptides each, but we have not validated this pooling scheme experimentally.

Lastly, we compared the experimental read out obtained by flow cytometry and fluorescent microscopy. Both methods yield consistent results with high correlation (*R*^2^ = 0.94 and *R*^2^ = 0.89, for MHCII-restricted and MHCI-restricted cell line, respectively; Figure 2F).

### Successful TCR deorphanization using *copepodTCR*

We next investigated whether a Spike peptide pool could be used to identify the epitope for a COVID-associated TCR with an unknown target. Lu and colleagues^17^ had previously reported paired TCR sequences for AIM+ cTfh cells after stimulation with a Spike peptide pool or recombinant Spike protein. We performed TCRdist analysis^18^ and picked two representative TCRs from the largest orphan cluster (labeled Lu_TCR1 and Lu_TCR2). Interestingly, a TCRbeta motif found in this cluster was also reported in ref.^19^ as a highly public, COVID-19-associated motif found in DRB1:16:01-positive and DQB1:05:01-positive donors. We cloned these TCRs into NFAT-GFP reporter Jurkat cell lines and using *copepodTCR* for CCP design and results interpretation found that the target epitope was the same for both: *S*_63−72_ (Figure 3A–D). We then confirmed this specificity by stimulating the reporter Jurkat cell lines with the individual peptide. Recognition was blocked by an anti-DR antibody and did not occur when PBMCs from a DRB1:16:01-negative, DQB1:05:01-positive donor were used as presenters, suggesting the peptide is presented by DRB1:16:01 and not DQB1:05:01.

**Figure 3.**
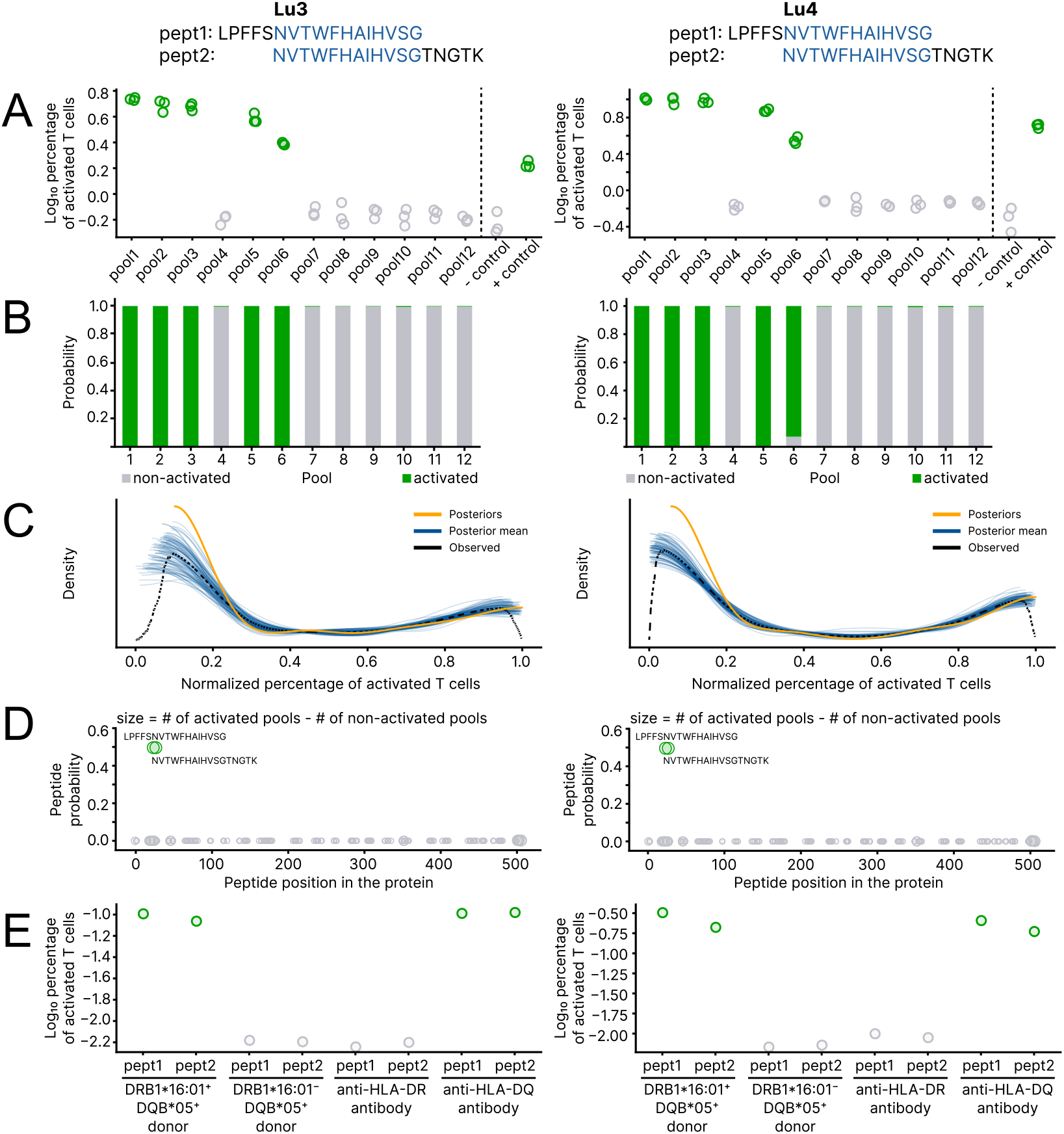
CopepodTCR design identifies specificity of TCRs with unknown targets. A. Flow cytometry read out as percentage of GFP-producing (activated) cells in the T cell activation assay (y-axis) for each pool in the design (x-axis), with pools detected as activated/non-activated in green and gray, respectively (as in B.). B. Inferred probability (from C) of each measurement coming from the distribution of activated pools (green) and from a distribution of non-activated pools (gray). C. Bayesian mixture model (blue and orange) of observed (black) pool activation signal for the read out obtained by flow cytometry. D. Per-peptide probabilities calculated from B, where the probability of one peptide equals the product of probabilities of pools to which it was added and 1 - probabilities of pools to which it was not added. The size of the circle shows the difference between the number of activated and non-activated pools to which a peptide was added. E. Both cell lines strongly recognize both individual overlapping peptides predicted from combinatorial pooling experiment. Recognition does not occur in presence of HLA-DR blocking antibody or if peptides are presented by DQB1:05:01-positive DRB1:16-negative donor.

## Discussion

We have demonstrated how *copepodTCR* can aid in the design and interpretation of CPP screens. Here, we discuss how we addressed design and results interpretation in distinct use case and outline remaining challenges in the design of peptide tiling screens and possible approaches to overcome them.

Tiling a protein sequence by overlapping peptides leads to the first and last peptide in the protein only sharing overlapping sequences with their successor or predecessor, respectively. If the TCR-specific epitope is located within these peptides, pools from only one address will be activated, in contrast to all other epitopes which activate the union of two addresses. In this edge case, the result will not be reported as a false negative, instead, *CopepodTCR* will alert the user that the activation scheme corresponds to an address associated with an end-position peptide.

Another limitation arises when dealing with peptide sets that do not have a consistent overlap length across all peptides. This inconsistency can lead to variable numbers of pools being activated, with more activated pools for increased epitope sharing across peptides and vice versa. Such inconsistency complicates the interpretation of experimental results, as these scenarios look indistinguishable from false positives or false negatives in the assay. However, *copepodTCR* contains an inconsistency check in the experimental setup, will alert the user of such issues and adjusts the interpretation of the results as addresses assigned to these peptides are known.

Potential peptide cross-reactivity poses a more significant challenge. In the case of cross-reactivity, several peptides containing similar epitopes would lead to the activation of the tested T cell. Cross-reactive epitopes often trigger different degrees of activation, which might aid in identifying these in post-hoc analyses. While cross-reactivity of TCRs in general is not a rare scenario, spanning orders of magnitude of different peptides that can be recognized by a single T cell^8^, it is still unlikely that two of these peptides will co-occur in a relatively small viral proteome.

For a large number of peptides to be tested, either the number of required pools or the number of peptides per pool will increase. For instance, testing 50,000 possible overlapping peptides would require a minimum of 35 pools, where each pool consists of approximately 5,700 peptides. For a stock peptide concentration of 10mM as we tested in this study, the individual peptide concentration after pooling in this set-up would be 1.75 *μ*M. At this low concentration, the hit rate of a T cell to encounter and thus get activated by its true cognate peptide is significantly decreased. To avoid this potential problem, the number of pools or the number of pools per peptide in the experimental setup can be adjusted to have a lower number of peptides per pool. However, from our dilution experiment (shown in Figure 2E), we were able to detect activation of the MHC class II restricted T cell line even with a 25-fold dilution for a pool with 253 peptides, corresponding to an individual peptide concentration per pool of 1.58 *μ*M, which is lower than the estimated concentration in the large screen exemplified above. Detecting activation of MHC class I restricted T cell lines required higher peptide concentrations. Peptides presented on MHC class I molecules are shorter than those presented on MHC class II molecules^20,21^. Prior to MHC loading, antigen-presenting cells use a multi-step process including proteases and transporter molecules to process the pooled peptides and achieve the required lengths. Differences in processing efficiency for peptides of short lengths might result in a weakened activation signal, lowering the detection efficiency for MHC class I restricted TCRs. In these cases, generating large peptide libraries by *in vitro* transcription and translation from oligonucleotides, as suggested by^22^, or from mini-gene libraries, as proposed by^23^, could be a cost-effective approach for the peptide library as input for the *copepodTCR*-designed CPP.

We used *copepodTCR* to identify a target epitope for the previously reported public COVID-19 associated TCR motif^17,19^ and found that these TCRs recognise the *S*_63−72_ epitope. Although reactivity to this region of Spike protein has been observed before^24,25^, no TCRs recognizing it have been reported. Here, we show that it is recognized by public TCR motifs. Interestingly, this epitope spans the *S*_69−70_ deletion whose frequency has fluctuated wildly over time (from 0% in July 2023 to 100% in July 2024 and 85% in July 2025). As of July 2025, most currently sequenced SARS-CoV-2 variants lack this epitope^26,27^. This makes *S*_63−72_-specific T cells an interesting model for studying differences in T cell immunity to repeated SARS-CoV-2 exposures.

In this work we validated the algorithm on MHC class I and II restricted cell lines with known specificity and used the algorithm to identify the cognate epitope of two known TCRs of thus far unknown specificity. With its straight-forward implementation and interpretation, our approach can scale to multiple cell lines, especially if the fraction of activated cells is measured by fluorescence microscopy. CPPs generated with *copepodTCR* could also be used for investigating the activation of primary cells, as is suggested for a subset of non-overlapping SARS-CoV-2 peptides in ref.^28^. In this protocol, sequencing of TCRs from T cells expressing activation surface markers after stimulation with peptide pools enables the identification of T cell clones co-occurring with peptide addresses, allowing identification of hundreds of TCR-peptide pairs in a single experiment. We expect that the small number of combinatorial pools combined with optimal assignment of peptides and robust error-correction enabled by the *copepodTCR* algorithm will be beneficial for this assay.

## Methods

### *copepodTCR* implementation

We developed a standalone Python package and graphical user interface for *copepodTCR*. The python package *copepodTCR* is available via *pip*. It relies on python packages codepub (version 2.3)^14^, pandas (version 1.5.3)^29^, NumPy (version 1.23.5)^30^, *Trimesh* (version 4.7.1)^31^, *manifold3d* (version 3.2.1)^32^, PyMC (version 5.25.1)^33^, Seaborn (version 0.12.2)^34^, Matplotlib (version 3.8.0)^35^, Plotly (version 6.2.0)^36^, and on Blender software (version 4.5.1)^37^. Its detailed documentation can be accessed at copepodTCR.readthedocs.io. The interactive user interface for *copepodTCR* (implemented with Shiny for Python (version 1.4.0)^38^) can be accessed at https://copepodtcr.cshl.edu/. Besides the *copepodTCR* and *codepub* Python packages, the tool uses Seaborn (version 0.12.2)^34^, pandas (version 2.3.1)^29^, Matplotlib (version 3.10.5)^35^, Plotly (version 6.3.0)^36^, and Biopython (version 1.85)^39^.

### Binary codes for designing combinatorial pooling schemes

*copepodTCR*’s implementation for generating a combinatorial pooling schemes builds on our software for the construction of balanced constant-weight gray codes for detecting consecutive positives (DCP-CWGCs)^14^: https://codepub.readthedocs.io/.

In brief, DCP-CWGCs for combinatorial peptide pooling propose a design scheme which consists of distinct binary vectors (binary addresses of peptides) that indicate the pools into which peptides are mixed. The binary addresses i) have a constant Hamming weight, and adjacent addresses have ii) a Hamming distance 2 and iii) unique OR-sums. These constraints enable the identification of consecutive positive items i.e. overlapping peptides sharing the same epitope, keep the number of tests on each item and on each pair of consecutive items constant, and facilitate error detection. To ensure stable and unbiased detection results across pools, we developed an implementation of DCP-CWGCs that produces *balanced* pooling arrangements, where the number of peptides per pool is approximately constant across all pools. Our implementation consists of a *branch-and-bound algorithm* (BBA) coupled with a recursive combination approach (rcBBA). The core algorithm, BBA, conducts a depth-first heuristic search for a path in the bipartite graph formed from the addresses of items and the unions of consecutive addresses. The rcBBA constructs long codes by recursive combination of several short, BBA-generated codes. Together, these implementations can construct balanced DCP-CWGCs for thousands of items in tractable time. Empirical run time analyses and pool balance, as well as mathematical descriptions of the algorithm, are provided in the Supplementary Note.

### Peptide mixture scheme

To aid with the experimental procedure, *copepodTCR* offers the creation of STL files, encoding the pipetting scheme necessary to create peptide pools based on the arrangement of experimental peptides in a 384-well plate. STL files are generated in Python using *Trimesh* (v4.7.1)^31^. Each card represents one pool, with holes positioned at the coordinates corresponding to the peptides designated for addition to that pool.

### Bayesian mixture model

To interpret the results of T cell activation assays, we developed a hierarchical Bayesian mixture model (Extended Data Fig 3). As input, the model requires an activation signal, *Activation*, expressed as a percentage of activated cells per replicate. We measured *Activation* by either flow cytometry or microscopy and computed the percentage of activated cells in one sample (flow cytometry) or the number of activated cells per sample divided by the total number of activated cells across all samples (microscopy). Regardless of the read out method, we normalized the activation per replicate by the maximal activation signal, so the highest activation value equals 1.

We assume that the *Activation* is composed of two distributions, representing activated and non-activated pools:

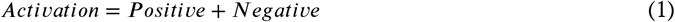

The non-activated (*Negative*) distribution is defined as truncated Normal with lower bound, *a*, and upper bound, *b*. The probability density function (PDF) of a truncated Normal distribution on the interval [*a, b*] is defined as the PDF of a standard normal distribution scaled such that the total probability over the interval [*a, b*] is 1.

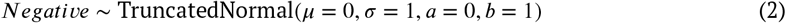

The parameters of the negative distribution are inferred either from the observed negative control (if measured) or from the pool with the lowest activation values:

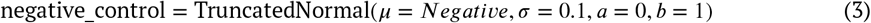

The activated (*Positive*) distribution is derived from the *Negative* distribution by adding a proportional *offset*. To ensure that the resulting distribution does not exceed 1, *offset* is defined as a proportion of the distance to the upper bound:

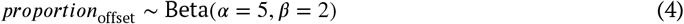

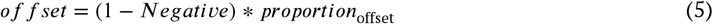

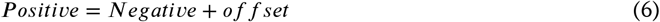

Each pool is assigned a value from a latent binary variable, *component*, that determines whether the pool was drawn from the *Positive* (= 0) or *Negative* distribution (= 1). This variable is modeled as a Bernoulli distribution parameterized by *probability*_*negative*_. *probability*_*negative*_ follows a Beta distribution whose prior is informed by the expected share of negative pools, which is obtained from the assay design. The expected share of negative pools is calculated as 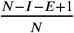, where *N* is the total number of pools, *I* is the peptide occurrence (i.e. the number of pools to which one peptide is added), and *E* is the number of peptides that share one epitope.

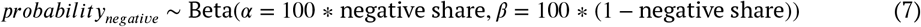

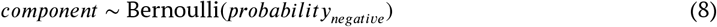

Each pool *i* has a pool-specific mean, *μ*_pool i_, and standard deviation, σ_pool i_, depending on its assigned *component*_*i*_ and defined using HalfNormal distributions. The PDF of a HalfNormal distribution is defined as the PDF of the standard Normal distribution with *μ* = 0 (restricted to non-negative values) and scaled so that the total probability over the interval [0, ∞] is 1.

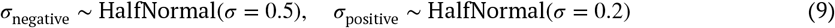

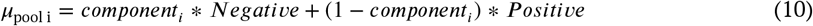

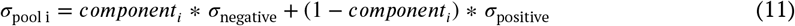

For each replicate *j*, associated with pool *i*, the observed normalized activation value is modeled using a truncated Normal distribution:

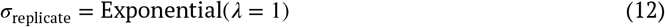

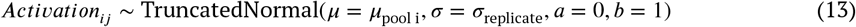

Posterior distributions are inferred using the No U-Turn Sampler [40]. The posterior mean of *component*_*i*_ for pool *i* quantifies the probability that pool *i* was drawn from a negative distribution (non-activated). A pool is considered non-activated if the posterior mean is equal to or greater than 0.5. Otherwise, the pool is considered to be activated.

The model was developed and fitted using PyMC (version 5.9.2)^33^.

Peptide probabilities were calculated as the product of probabilities of pools to which they were added and 1 - the product of probabilities of pools to which they were not added.

### *In silico* combinatorial pooling experiments

The *in silico* data generation process consisted of four steps (Extended Data Fig 4A).

1. Generation of peptides. {100 or 1000} peptides were generated using a sliding window approach from {1 or 10} random amino acid sequences (proteins), with a shift of {4 or 6} amino acids and a peptide length of {14 or 18} amino acids.
2. Peptide pooling. Peptides were assigned to {10, 12, or 14} pools using the *codepub* pooling algorithm. For each tested number of peptides and number of pools, appropriate peptide occurrence was selected such that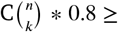 number of peptides, and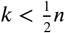, where *n* is the number of pools, and *k* is the peptide occurrence.
3. Cognate peptides selection. From the set of all epitopes present in the peptide library, an epitope of length {8 or 14} amino acids was randomly chosen. Peptides sharing the selected epitope were designated as cognate peptides.
4. Activation signal generation (Extended Data Fig 4B). Pools containing cognate peptides were labeled as activated. All pools without cognate peptides were labeled non-activated.

Each pool was simulated with {1, 2, or 3} replicate measurements. Additionally, we tested for the effect of experimental errors by introducing false negatives: {0, 1} randomly selected activated pools were reclassified and sampled from the *Negative* distribution.

The activation signal represents the percentage of activated T cells, and therefore was constrained to the range from 0 to 100; thus in simulation, activation values were lower and upper bound by 0 and 100.

To simulate this activation signal, we used the scaled sum of two truncated normal distributions, reflecting the negative and positive pool signals. The *Negative* distribution was defined as Truncated Normal with mean = {5, 15, or 45}, standard deviation = 3, and lower and upper bounds equal to 0 and 100, respectively:

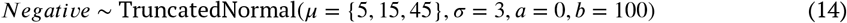

The *Positive* distribution was defined deterministically from *Negative* by adding a *signal* distribution. The *signal* followed a TruncatedNormal with mean = {5, 15, or 45}, standard deviation = 3, and lower and upper bounds equal to 0 and 100:

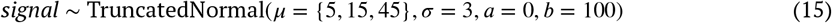

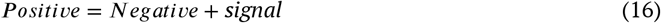

Out of all pools containing a single cognate peptide, one pool was randomly selected to exhibit a reduced activation signal, introduced to reflect the effect of different peptide contexts on epitope processing, loading, and recognition. The activation values of this pool were sampled from a scaled version of the positive distribution by multiplying the *Positive* by a scaling factor = {0.2 or 0.8}:

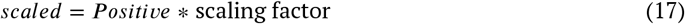

Pools were simulated from the corresponding distribution, and activation values for *r* = {1, 2, or 3} replicates were sampled from a corresponding pool.

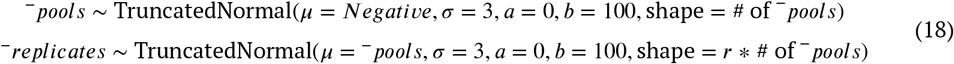

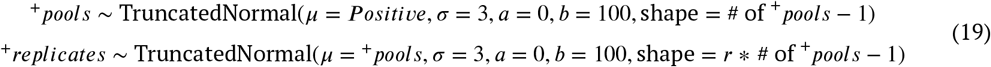

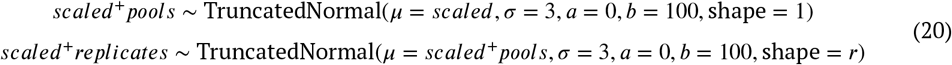

Negative control measurements were drawn separately from the *Negative* distribution:

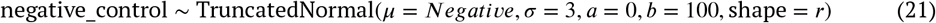

To obtain negative data, i.e., without any activation signal, all pools were considered negative, and consequently, activation values for all pools were sampled from the *Negative* distribution.

The model was sampled using the No-U-Turn Sampler (NUTS) [40] with a single draw, as its purpose was to simulate from the prior predictive distribution, not to perform inference. Posterior means were computed across the chain dimension for each pool replicate.

In total, we tested all combinations of the parameters described above, resulting in 19,440 independent simulations, including 6,480 negative data (without activation signal) simulations.

To test the model using the data with more diverse expected shares of negative pools, we used the same parameters except for the number of pools = {10, 15, 20}, number of peptides = {90, 100, 500, 1000}, number of proteins = 1, overlap length = 4, peptide length = 14, epitope length = 8. Peptide occurrence was selected such that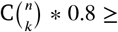 number of peptides, and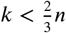, where *n* is the number of pools, and *k* is the peptide occurrence. This set of parameters resulted in 12,636 independent simulations, including 4,212 negative data simulations.

### Peptide pooling

Individual 17-mer peptides, overlapping by 12 amino acids from the SARS-CoV-2 Spike protein (Uniprot acc. P0DTC2, synthesized by Mimotopes), were diluted to 10mM in a matching solvent according to the manufacturer’s solubility test. The diluted peptides were transferred to a 384-well plate (951020702, Eppendorf).

For the dilution experiment, we mixed all 253 peptides in a single pool and then diluted the mixture 5- and 25-fold, corresponding to concentrations for each individual peptide in these pools of 39.5 *μ*M, 7.91 *μ*M, and 1.58 *μ*M, respectively.

We used *copepodTCR* to design a peptide pooling scheme and corresponding STL files for punched cards matching an empty 12.5 *μ*l GripTIPS box (Integra), generated G-code with Cura v. 5.1.0 (Ultimaker), and printed them with PLA on a 3D printer (Ender V3, Creality). Each punched card encoded the pooling strategy for a single combinatorial pool: we placed a card on top of an empty tip box, filled open holes with tips, and then use this patterned pipette tip array to transfer 2.5 *μ*l from the source plate to the lid of the tip box using a ViaFLO 384-channel electronic pipette (Integra). The tip box lid was then pulse centrifuged at 1000g to collect droplets in the corner and pooled peptides were transferred into an 8-tube strip and stored at -80^°^C before use.

### Stimulation of reporter T cell lines with combinatorial pools

For validation of our approach, we used a NFAT-GFP reporter Jurkat 76.7 cell lines from ref.^41^ (YLQ TCR 1) and ref.^15^ (TCR6.3), both with known specificity. Jurkat cells (10^5^) were co-cultured in a round-bottom 96-well cell culture plate (Corning 3799) in 100 *μ*l RPMI-1640 media (Gibco) supplemented with 10% FBS, 1% penicillin-streptomycin, containing 1 *μ*g/ml anti-CD28 antibody (BD Biosciences, 555725) and 1 *μ*g/ml of anti-CD49d antibody (BD Biosciences, 555501), with 10^5^ PBMCs from a healthy HLA-A02+ and HLA-DPB1*04:02+ donor (matching the MHC specificity of the YLQ TCR 1 and TCR6.3, respectively) pulsed with 1 *μ*l of the given combinatorial pool for 16 hours. For positive controls we used PMA/Ionomycin cocktail (Invitrogen, 00-4970-93) according to manufacturer protocol, or single YLQPRTFLL or CTFEYVSQPFLMDLEGK peptide (known YLQ TCR 1 TCR6.3 target epitopes respectively) at final concentration in media 1*μ*M. Each co-culture experiment was performed in triplicates. Plates were washed with 300 *μ*l of FACS buffer (DPBS with 0.05% BSA and 2mM EDTA) and resuspended in 300 *μ*l of FACS buffer. Subsequently, 150 *μ*l were transferred into a 96-well flat-bottom plate (Corning 3596) and acquired on the Incucyte S3 (Sartorius) with a 4x objective. Green object count for each well was determined using Incucyte Analysis Software with Surface Fit segmentation, no edge split, 2.0 GCU Threshold and *<* 750*μm*^2^ Area filter. Simultaneously, a second plate was acquired on BD Symphony A2 with a high throughput sampler (Extended Data Fig 6 for gating strategy). Flow cytometry data was analysed with FlowJo (v. 10.8.1). Results were interpreted using the Bayesian Mixture model of *copepodTCR*.

### Deorphanization of patient-derived public TCRs

To select T cell motifs for deorphanization, we used single-cell TCR sequencing data from activated T cells stimulated with inactivated virus, recombinant spike protein, or S, M, and N peptide pools. After grouping TCR sequences with TCRdist ≤ 120, we found that the largest cluster corresponded to the previously characterized TRAV35/TRAJ42 TCR motif^15^ recognizing the CTFEYVSQPFLMDLEGK epitope (similar to TCR6.3). We next selected two representative TCRs from the second largest cluster, Lu1: TCR*α* TRAV5, CDR3: CAESNNDMRF, TRAJ43; TCR*β* TRBV12-4, CDR3: CASSRTGRGSSYNSPLHF, TRBJ1-6; and Lu2: TCR*α* TRAV23, CDR3: CAGAEGGKLIF, TRAJ23; TCR*β* TRBV12-4, CDR3: CASSRTGFRSSYNSPLHF, TRBJ1-6.

The *αβ* TCR genes were synthesized (GenScript) and transduced into NFAT-GFP reporter Jurkat 76.7 cells as described previously^15^. We next performed a co-culture experiment with combinatorial spike pools as described above. To validate the results, we set up another co-culture experiment with individual peptides at 1 *μ*M, with and without anti-HLA-DR (BioLegend, 307666; 5 *μ*g/mL) or anti-HLA-DQ (Novus Biologicals, NBP2-45041; 1 *μ*g/mL) blocking antibodies at final concentration.

## Author contributions

VAK, PGT, MVP, and HVM conceptualized the work; VAK developed the software with support from GH and implemented the user interface together with SRC; CB and QH advised on algorithm design; VAK, DJP and HVM developed the hierarchical model; VAK conducted the formal analyses; MVP, AAM, KAR, AJS and VAK performed the experiments; VAK, MVP, and HVM wrote the original draft; all authors reviewed & edited the final draft; MVP, PGT, and HVM supervised the work.

## Code availability

Custom analysis code was written in python (version ≥ 3.10.11). Code and data to re-produce all figures in the manuscript are freely available on GitHub: https://github.com/meyer-lab-cshl/copepodTCR_paper.git.

## Competing interests

PGT has consulted for Pfizer, JNJ, Cytoagents, 10X, and Illumina, and serves on the SAB for Shennon Bio and Immunoscape. PGT, AAM, and MVP have patents related to TCR amplification, cloning, and applications thereof. The authors declare no other competing interests.

### Acknowledgments

The authors thank Jeremy Shaw, Beiyun Liu, Luigi Mari and R.K. Subbarao Malireddi for their help with microscopy assays; Katherine Richards and Andrea Sant for discussions on epitope discovery and peptide libraries; and Anastasia Troshina for the *copepodTCR* logo design.

---

**Extended Data Figure 1.**
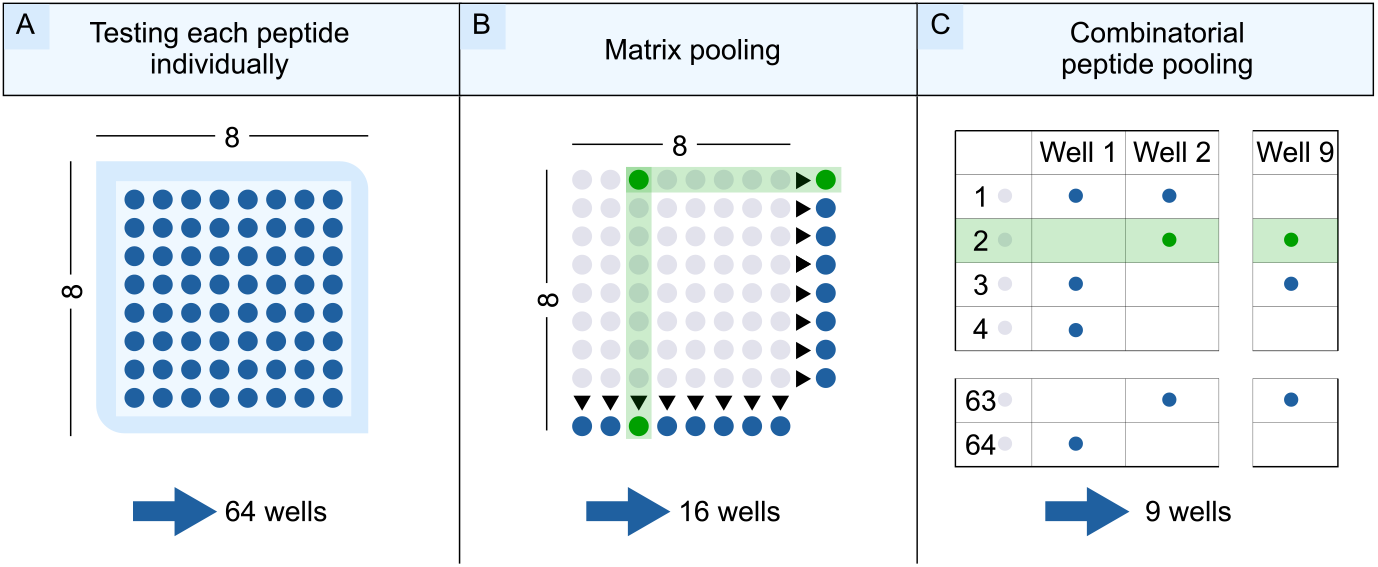
Experimental layout of TCR peptide screen. Three common approaches to designing a TCR peptide screen. They differ in experimental complexity (high to low, left to right) and computational design (low to high, left to right). Details are described in the main text.

**Extended Data Figure 2.**
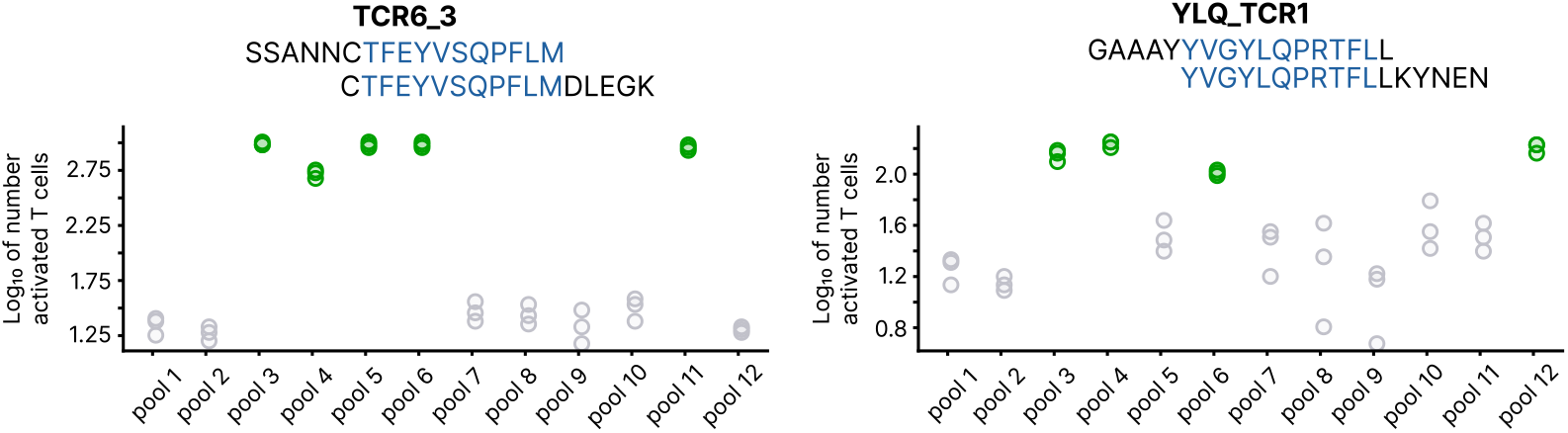
Fluorescent microscopy read out. Percentage of GFP-producing (activated) cells in the T cell activation assay (y-axis) for each pool in the design (x-axis), with pools detected as activated/non-activated in green and grey, respectively. Results shown for the T cell reporter cell lines expressing DPB1*04:(01/02)-restricted TCR6.3 (left column) and HLA-A02+ restricted YLQ-TCR1 (right column) with their known, cognate peptides indicated in blue letters in the peptide sequence.

**Extended Data Figure 3.**
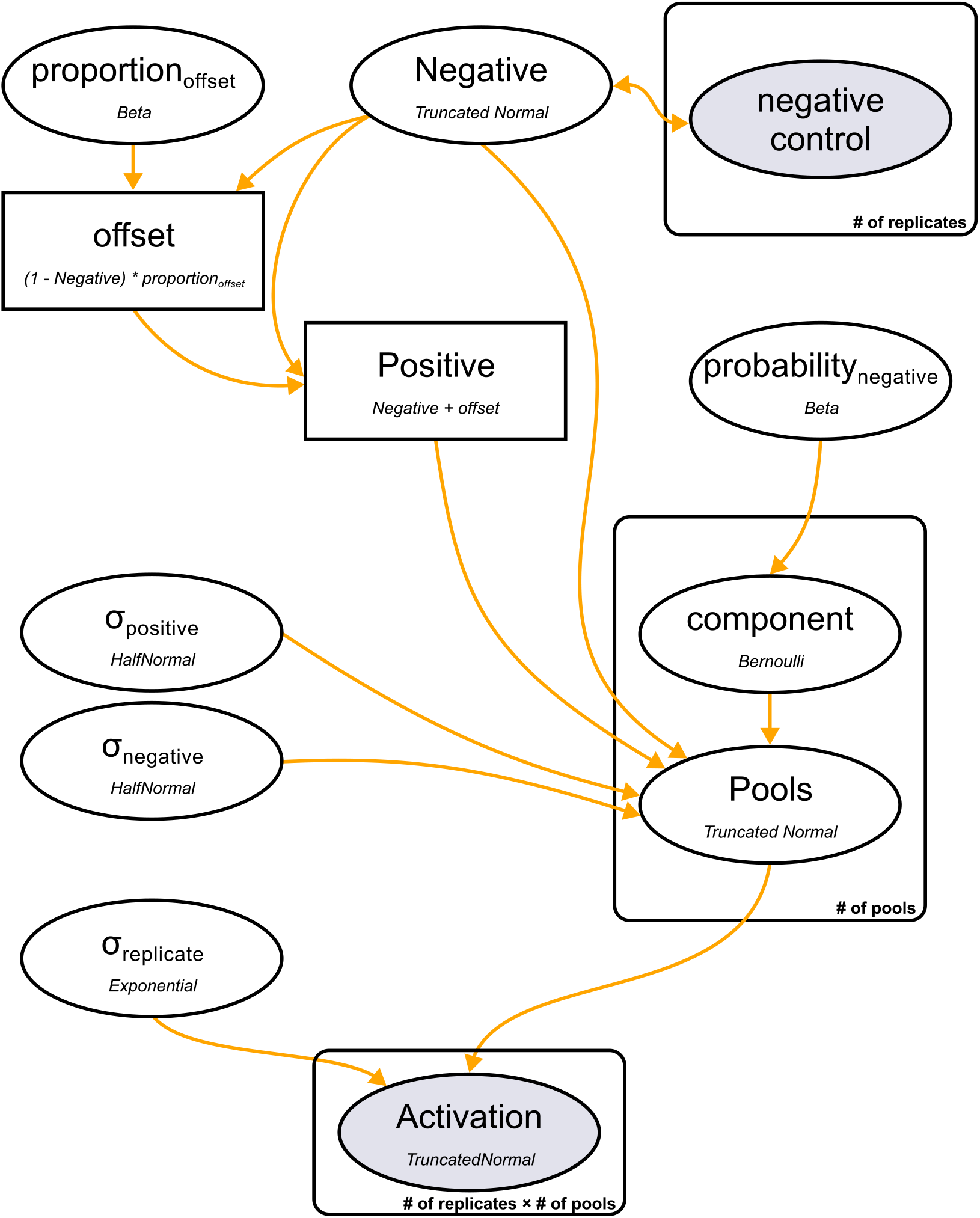
Bayesian Mixture model of activation signal. Plate notation of the model. The activation signal *Activation* is modeled as a mixture of the *Negative* and *Positive* distributions, both truncated with lower and upper bounds fixed at 0 and 1, respectively. The *Negative* distribution follows a TruncatedNormal distribution, with its location inferred from the negative control measurements. The *Positive* distribution is derived by adding an *offset* to the *Negative* distribution. The *offset* is learned as a proportion of the remaining distance to the upper bound. Each pool is assigned a latent binary variable *component*, which follows a Bernoulli distribution parameterized by *probability*_*negative*_. This probability is drawn from a Beta distribution reflecting the expected fraction of negative pools. Each pool *i* has a pool-specific mean and standard deviation, conditional on its assigned *component*_*i*_. For each replicate *j*, associated with pool *i*, the observed normalized activation value is modeled using a TruncatedNormal distribution.

**Extended Data Figure 4.**
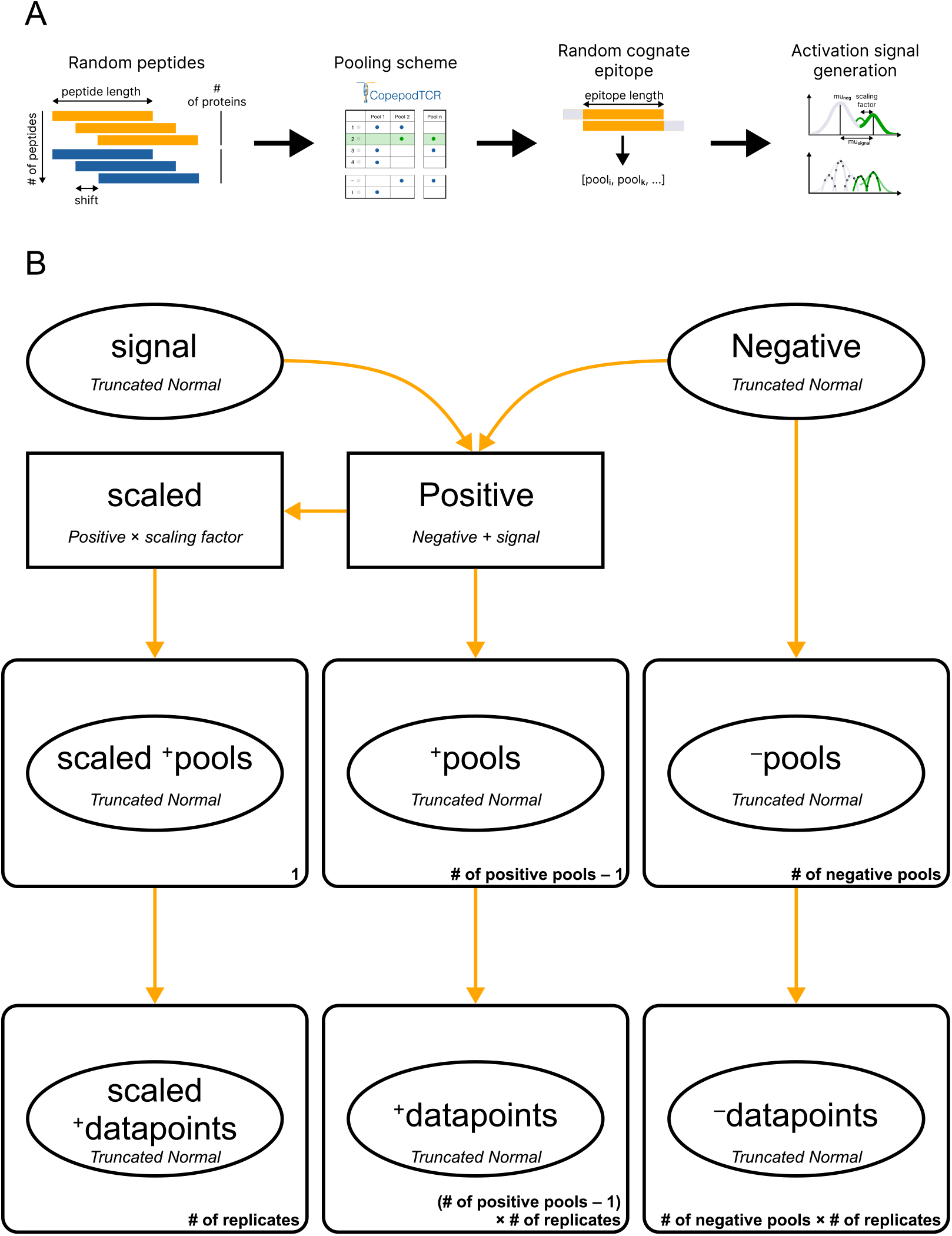
*In silico* data generation. **A**. Schematic of *in silico* data generation. Peptides are generated from random amino acid sequences (proteins) using a sliding window approach with a defined shift and peptide lengths. A peptide pooling scheme is then constructed using a specified number of pools and peptide occurrence. From all possible epitopes of a defined length present in the tested peptides, a random epitope is selected, and peptides containing the selected epitope are designated as cognate. Based on the pooling scheme, pools containing cognate peptides are labeled as activated. Activation values for all pools are subsequently simulated using the model described in panel B. **B**. Plate notation of the simulation. Activation is simulated using TruncatedNormal distributions with pool-specific parameters. *μ*_pool_ is drawn from one of three distributions: *Positive, Scaled Positive*, or *Negative*. The *Negative* distribution follows a TruncatedNormal distribution. The *Positive* distribution is derived deterministically by adding a *signal* distribution to the *Negative*, where *signal* follows its own TruncatedNormal distribution. For one of the activated pools, the activation signal is drawn from a *Scaled Positive* distribution defined as *Positive* multiplied by a fixed scaling factor.

**Extended Data Figure 5.**
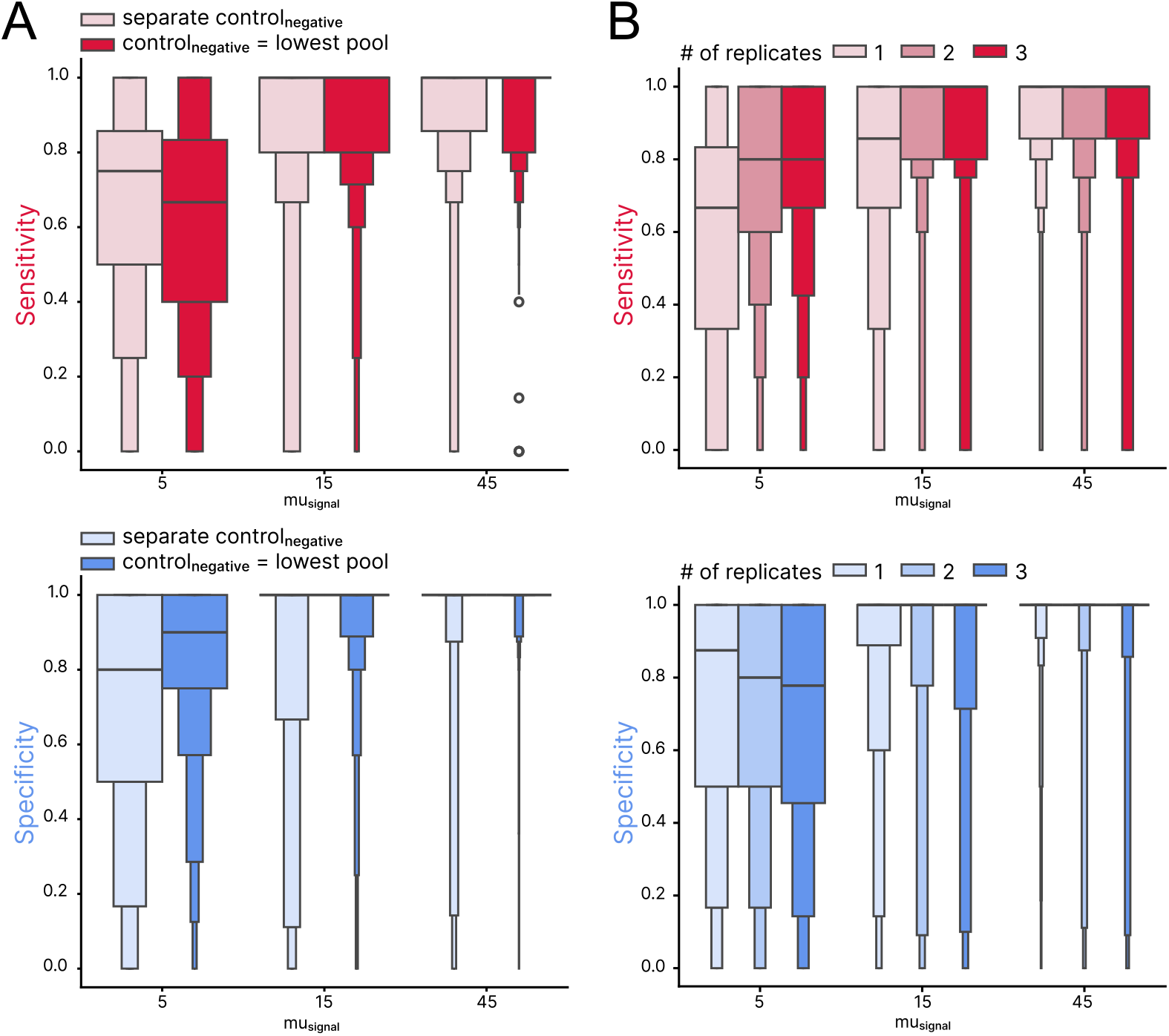
Activation model predictions on synthetic data. A. Sensitivity and specificity of the activation model with parameters for negative distribution inferred from separate negative control data (separate control_negative_) or from the pool with the lowest activation signal (control_negative_ = lowest pool). The model’s performance in both scenarios depends on signal intensity (*mu*_*signal*_, representing the difference in percentage of activated cells between positive and negative distributions), where in simulations with a weak signal (*mu*_*signal*_ = 5), the sensitivity of the model is higher with a separate negative control, but specificity is lower. B. Sensitivity and specificity of the activation model are similar across simulations with different numbers of replicates per pool (1, 2, or 3 replicates). Sensitivity slightly increases with the number of replicates per pool (upper panel), but specificity decreases (lower panel), especially with weak signal (lower panel, *mu*_*signal*_ = 5.)

**Extended Data Figure 6.**
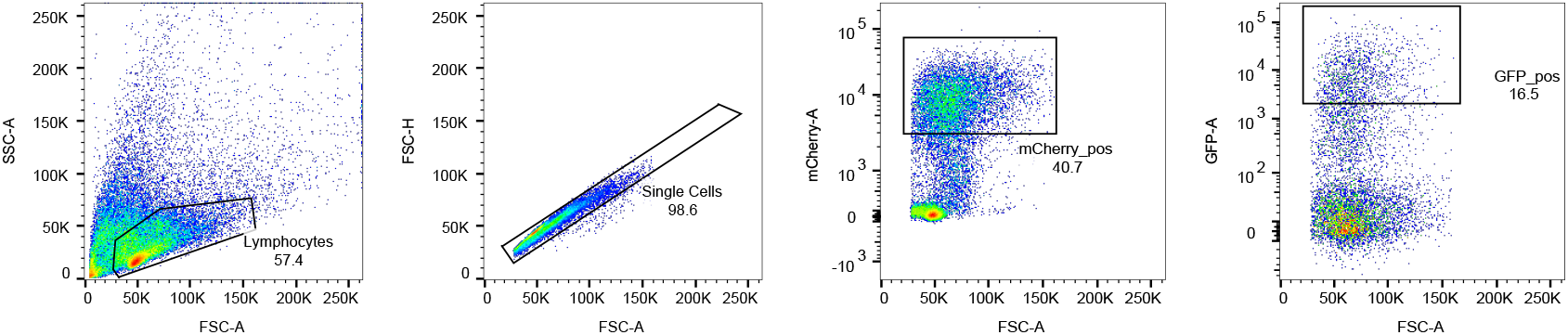
Gating strategy to identify activated NFAT-GFP reporter TCR6.3 Jurkat cells. Activated Jurkat cells are mCherry+/GFP+. mCherry is constitutively expressed on all cells expressing TCR6.3. GFP is under NFAT control and expressed after TCR engagement.

## References

1. Sbai, H, Mehta, A & DeGroot, A. Use of T cell epitopes for vaccine development. Current drug targets-Infectious disorders 1. doi:10.2174/1568005014605955 (2001).

2. Waldman, AD, Fritz, J. & Lenardo, MJ. A guide to cancer immunotherapy: from T cell basic science to clinical practice. Nature Reviews Immunology 20. doi:10.1038/s41577-020-0306-5 (2020).

3. Prinz, JC. Immunogenic self-peptides-the great unknowns in autoimmunity: Identifying T-cell epitopes driving the autoimmune response in autoimmune diseases. Frontiers in Immunology 13. doi:10.3389/fimmu.2022.1097871 (2023).

4. Davis, M. & Bjorkman, PJ. T-cell antigen receptor genes and T-cell recognition. Nature 334. doi:10.1038/334395a0 (1988).

5. Zhao, W & Sher, X. Systematically benchmarking peptide-MHC binding predictors: From synthetic to naturally processed epitopes. PLOS Computational Biology 14. doi:10.1371/journal.pcbi.1006457 (2018).

6. Meysman, P, Barton, J, Bravi, B, Cohen-Lavi, L, Karnaukhov, V, Lilleskov, E, Montemurro, A, Nielsen, M, Mora, T, Pereira, P, et al. Benchmarking solutions to the T-cell receptor epitope prediction problem: IMMREP22 workshop report. ImmunoInformatics 9. doi:10.1016/j.immuno.2023.100024 (2023).

7. Dens, C, Laukens, K, Bittremieux, W & Meysman, P. The pitfalls of negative data bias for the T-cell epitope specificity challenge. Nature Machine Intelligence 5. doi:10.1038/s42256-023-00727-0 (2023).

8. Banerjee, A, Pattinson, DJ, Wincek, CL, Bunk, P, Axhemi, A, Chapin, SR, Navlakha, S & Meyer, HV. T cell receptor cross-reactivity prediction improved by a comprehensive mutational scan database. Cell Systems. doi:10.1016/j.cels.2025.101345 (2025).

9. Chaves, FA, Lee, AH, Nayak, JL, Richards, K. & Sant, AJ. The Utility and Limitations of Current Web-Available Algorithms To Predict Peptides Recognized by CD4 T Cells in Response to Pathogen Infection. The Journal of Immunology 188. doi:10.4049/jimmunol.1103640 (2012).

10. Joglekar, A. & Li, G. T cell antigen discovery. Nature methods 18. doi:10.1038/s41592-020-0867-z (2021).

11. Fiore-Gartland, A, Manso, BA, Friedrich, DP, Gabriel, EE, Finak, G, Moodie, Z, Hertz, T, De Rosa, SC, Frahm, N, Gilbert, PB, et al. Pooled-peptide epitope mapping strategies are efficient and highly sensitive: an evaluation of methods for identifying human T cell epitope specificities in large-scale HIV vaccine efficacy trials. PloS one 11. doi:10.1371/journal.pone.0147812 (2016).

12. Klinger, M, Pepin, F, Wilkins, J, Asbury, T, Wittkop, T, Zheng, J, Moorhead, M & Faham, M. Multiplex identification of antigen-specific T cell receptors using a combination of immune assays and immune receptor sequencing. PloS one 10. doi:10.1371/journal.pone.0141561 (2015).

13. Snyder, TM, Gittelman, RM, Klinger, M, May, DH, Osborne, EJ, Taniguchi, R, Zahid, HJ, Kaplan, IM, Dines, JN, Noakes, MT, et al. Magnitude and Dynamics of the T-Cell Response to SARS-CoV-2 Infection at Both Individual and Population Levels. doi:10.1101/2020.07.31.20165647 (2020).

14. He, G, Kovaleva, VA, Barton, C, Thomas, PG, Pogorelyy, MV, Meyer, H. & Huang, Q. Unbiased and Error-Detecting Combinatorial Pooling Experiments with Balanced Constant-Weight Gray Codes for Consecutive Positives Detection. doi:10.48550/arXiv.2502.08214 (2025).

15. Mudd, PA, Minervina, AA, Pogorelyy, MV, Turner, JS, Kim, W, Kalaidina, E, Petersen, J, Schmitz, AJ, Lei, T, Haile, A, et al. SARS-CoV-2 mRNA vaccination elicits a robust and persistent T follicular helper cell response in humans. Cell 185. doi:0.1016/j.cell.2021.12.026 (2022).

16. Shomuradova, AS, Vagida, MS, Sheetikov, SA, Zornikova, KV, Kiryukhin, D, Titov, A, Peshkova, IO, Khmelevskaya, A, Dianov, DV, Malasheva, M, et al. SARS-CoV-2 Epitopes Are Recognized by a Public and Diverse Repertoire of Human T Cell Receptors. Immunity 53. doi:10.1016/j.immuni.2020.11.004 (2020).

17. Lu, X, Hosono, Y, Nagae, M, Ishizuka, S, Ishikawa, E, Motooka, D, Ozaki, Y, Sax, N, Maeda, Y, Kato, Y, et al. Identification of conserved SARS-CoV-2 spike epitopes that expand public cTfh clonotypes in mild COVID-19 patients. Journal of Experimental Medicine 218. doi:10.1084/jem.20211327 (2021).

18. Mayer-Blackwell, K, Schattgen, S, Cohen-Lavi, L, Crawford, JC, Souquette, A, Gaevert, JA, Hertz, T, Thomas, PG, Bradley, P & Fiore-Gartland, A. TCR meta-clonotypes for biomarker discovery with tcrdist3 enabled identification of public, HLA-restricted clusters of SARS-CoV-2 TCRs. eLife 10. doi:10.7554/eLife.68605 (2021).

19. Vlasova, EK, Nekrasova, AI, Komkov, AY, Izraelson, M, Snigir, EA, Mitrofanov, SI, Yudin, VS, Makarov, VV, Keskinov, AA, Korneeva, D, et al. Robust detection of SARS-CoV-2 exposure in the population using T-cell repertoire profiling. doi:10.1101/2023.11.08.566227 (2023).

20. Szeto, C, Lobos, CA, Nguyen, A. & Gras, S. TCR recognition of peptide–MHC-I: Rule makers and breakers. International journal of molecular sciences 22. doi:10.3390/ijms22010068 (2020).

21. La Gruta, NL, Gras, S, Daley, SR, Thomas, P. & Rossjohn, J. Understanding the drivers of MHC restriction of T cell receptors. Nature Reviews Immunology 18. doi:10.1038/s41577-018-0007-5 (2018).

22. Zhang, SQ, Ma, KY, Schonnesen, AA, Zhang, M, He, C, Sun, E, Williams, CM, Jia, W & Jiang, N. High-throughput determination of the antigen specificities of T cell receptors in single cells. Nature Biotechnology. doi:10.1038/nbt.4282 (2019).

23. Cattaneo, CM, Battaglia, T, Urbanus, J, Moravec, Z, Voogd, R, De Groot, R, Hartemink, KJ, Haanen, JBAG, Voest, EE, Schumacher, TN, et al. Identification of patient-specific CD4+ and CD8+ T cell neoantigens through HLA-unbiased genetic screens. Nature Biotechnology 41. doi:10.1038/s41587-022-01547-0 (2023).

24. Karsten, H, Cords, L, Westphal, T, Knapp, M, Brehm, TT, Hermanussen, L, Omansen, TF, Schmiedel, S, Woost, R, Ditt, V, et al. High-resolution analysis of individual spike peptide-specific CD4+ T-cell responses in vaccine recipients and COVID-19 patients. Clinical & Translational Immunology 11. doi:10.1002/cti2.1410 (2022).

25. Keller, MD, Harris, KM, Jensen-Wachspress, MA, Kankate, VV, Lang, H, Lazarski, CA, Durkee-Shock, J, Lee, PH, Chaudhry, K, Webber, K, et al. SARS-CoV-2–specific T cells are rapidly expanded for therapeutic use and target conserved regions of the membrane protein. Blood 136. doi:10.1182/blood.2020008488 (2020).

26. Hadfield, J, Megill, C, Bell, SM, Huddleston, J, Potter, B, Callender, C, Sagulenko, P, Bedford, T & Neher, RA. Nextstrain: real-time tracking of pathogen evolution. Bioinformatics 34. doi:10.1093/bioinformatics/bty407 (2018).

27. Khare, S, GISAID Global Data Science Initiative (GISAID), Munich, Germany, Gurry, C, Freitas, L B, Schultz, M, Bach, G, Diallo, A, Akite, N, Ho, J, Tc Lee, R. et al. GISAID’s Role in Pandemic Response. China CDC Weekly 3. doi:10.46234/ccdcw2021.255 (2021).

28. Snyder, TM, Gittelman, RM, Klinger, M, May, DH, Osborne, EJ, Taniguchi, R, Zahid, HJ, Kaplan, IM, Dines, JN, Noakes, MT, et al. Magnitude and dynamics of the T-cell response to SARS-CoV-2 infection at both individual and population levels. MedRxiv. doi:10.1101/2020.07.31.20165647 (2020).

29. McKinney, W. Data Structures for Statistical Computing in Python in Proceedings of the 9th Python in Science Conference (2010). doi:10.25080/Majora-92bf1922-00a.

30. Harris, CR, Millman, KJ, van der Walt, SJ, Gommers, R, Virtanen, P, Cournapeau, D, Wieser, E, Taylor, J, Berg, S, Smith, NJ, et al. Array programming with NumPy. Nature 585. doi:10.1038/s41586-020-2649-2 (2020).

31. Dawson-Haggerty et al. trimesh version 3.2.0. https://trimsh.org/.

32. Lalish, E. manifold3d version 3.2.1. 2025. https://pypi.org/project/manifold3d/.

33. Oriol, AP, Virgile, A, Colin, C, Larry, D J.,, FC, Maxim, K, Ravin, K, Jupeng, LC, LC, A., MO, et al. PyMC: A Modern and Comprehensive Probabilistic Programming Framework in Python. PeerJ Computer Science 9. doi:10.7717/peerj-cs.1516 (2023).

34. Waskom, ML. seaborn: statistical data visualization. Journal of Open Source Software 6. doi:10.21105/joss.03021 (2021).

35. Hunter, JD. Matplotlib: A 2D graphics environment. Computing in Science & Engineering 9. doi:10.1109/MCSE.2007.55 (2007).

36. Inc., PT. Collaborative data science

37. Blender Foundation and Community. Blender version 4.5.1. 2025. https://www.blender.org/.

38. Development team, TS. Shiny for Python https://github.com/rstudio/py-shiny.

39. Cock, PJ, Antao, T, Chang, JT, Chapman, BA, Cox, CJ, Dalke, A, Friedberg, I, Hamelryck, T, Kauff, F, Wilczynski, B, et al. Biopython: freely available Python tools for computational molecular biology and bioinformatics. Bioinformatics 25 (2009).

40. Hoffman, MD, Gelman, A, et al. The No-U-Turn sampler: adaptively setting path lengths in Hamiltonian Monte Carlo. J. Mach. Learn. Res. 15. doi:10.5555/2627435.2638586 (2014).

41. Minervina, AA, Pogorelyy, MV, Kirk, AM, Crawford, JC, Allen, EK, Chou, CH, Mettelman, RC, Allison, KJ, Lin, CY, Brice, DC, et al. SARS-CoV-2 antigen exposure history shapes phenotypes and specificity of memory CD8+ T cells. Nature Immunology 23. doi:10.1038/s41590-022-01184-4 (2022).

