## Supplementary Notes for "Identification of Antigen-Specific T Cell Receptors with Combinatorial Peptide Pooling"

[1]Simons Center for Quantitative Biology, Cold Spring Harbor Laboratory, Cold Spring Harbor, NY 11724, USA; [2]Department of Pathobiological Sciences, School of Veterinary Medicine, University of Wisconsin-Madison, Madison, WI 53711, USA; [3]School of Electronic and Information Engineering, Beihang University, Beijing 100191, China; [4]Birkbeck, University of London, WC1E 7HX London, UK; [5]Department of Host-Microbe Interaction, St. Jude Children's Research Hospital, Memphis, TN 38105, USA

### Supplementary Notes

#### Algorithmic constraints for the combinatorial peptide pooling scheme

Let's consider a single protein $E$. We create a library of peptides by a sliding window approach across its sequence, i.e., each peptide overlaps with its predecessor and successor by the constant number of amino acids. The only exception is the first and last peptide in the protein, which only overlap with the successor and predecessor respectively. In our experiment, we have a total number of $M$ overlapping peptides $E_j$ derived from $E$:

$$E = \{E_1, \ldots, E_M\}, \text{ with } overlap(E_j, E_{j+1}) = const \tag{1}$$

for which we want to find a mixing scheme, into $n$ pools $p$:

$$p = \{p_1, \ldots, p_n.\} \tag{2}$$

The algorithm needs to find the optimal distribution of peptides into the pools, such that the number of pools $n$ is minimized, the total occurrence of each peptide across pools equals $x$, where $x$ is minimized, and the total number of peptides per pool is approximately constant.

We consider the distribution of peptides $E_j$ into pools $p_i$ as assigning addresses $a_j$ to each peptide $E_j$. An address $a_j$ is represented by a binary string with $n$ number of digits, digit $b_i$ encoding the presence of a peptide $E_j$ in pool $p_i$. The number of digits equal to 1 in any address $a_j$ should equal $x$:

$$\forall j : \sum_{i=1}^{n} b_{ji} = x \tag{3}$$

The construction of the pools is limited by the combination of the following constraints:

- The number of peptides per pool should be approximately the same for each pool:

$$\overline{\overline{p_i}} \approx w \text{ where } w = \frac{M * x}{n} \tag{4}$$

- Each address $a_j$ differs only in one pool from its successor $a_{j+1}$:

$$a_{j+1} \text{ with } D_H(a_j, a_{j+1}) = 2 \tag{5}$$

  where $D_H$ is the Hamming distance.
- For all other addresses, the Hamming distance is equal or greater than 2:

$$D_H(a_j, a_k) \geq 2 \text{ where } |j - k| \geq 2 \tag{6}$$

- The Hamming distance between the union of two adjacent addresses and any other union of adjacent addresses is equal or greater than 2:

$$\forall j, k \,:\, D_H\left(a_j \cup a_{j+1},\, a_k \cup a_{k+1}\right) \geq 2 \tag{7}$$

### Efficient construction of combinatorial pooling schemes for high-throughput experimental assays.

We have developed two algorithms for constructing of balanced constant-weight Gray codes for detecting consecutive positives (DCP-CWGCs)[1] with varying parameters (Supplementary Fig 1A). The first algorithm, *branch-and-bound algorithm* (BBA), encodes the position of each peptide by a binary sequence (address), where each bit represents a pool, and "1" means that a peptide was added to that pool and "0" that it was not. According to constraint Eq. (3), the weight of the code (number of "1" in any given address) remains constant in the experimental setup and represents a peptide occurrence, i.e. the number of pools to which one peptide was added. BBA then performs a search for a path in the bipartite graph whose nodes represent individual addresses and unions of two adjacent addresses. The algorithm prioritizes balance by ensuring that each pool (i.e., bit position) is utilized approximately equally across the entire scheme.

To improve scalability, particularly for codes of moderate to long length, we developed a method based on *recursive combination together with branch-and-bound algorithm* (rcBBA). This approach constructs long DCP-CWGCs by recursively combining several shorter codes generated by BBA. For moderate and long codes, rcBBA exhibits substantially decreased running time while maintaining a near-perfect balance. We implemented both BBA and rcBBA in the python package *codepub*, and algorithm statistics presented below are based on this implementation.

### *copepodTCR* demonstrates consistent and fast performance for CPP assays

*copepodTCR* can design complex peptide pooling libraries in less than a minute. For instance, to design a peptide pooling scheme with 1,000 overlapping peptides distributed over 16 pools and each peptide added to 4 pools takes less than 3 seconds (Supplementary Fig 1B).

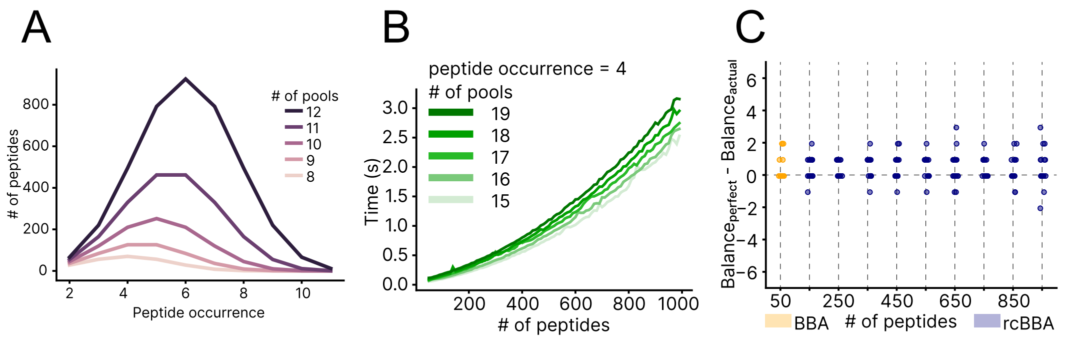


**Supplementary Figure 1. Algorithm statistics.** A. Interaction between the number of peptides, the number of pools, and peptide occurrence parameters in the CPP. B. Empirical runtime analyses showing that the *codepub* rcBBA algorithm depends on the number of pools and peptides (displayed for constant number of pools per peptide = 4). C. Deviation of *codepub*-derived experimental balance ($Balance_{actual}$) from optimal balance ($Balance_{perfect}$). The difference is displayed for a simulation of 16 pools and a constant number of pools per peptide = 4. For the number of peptides $\leq$ 50, the BBA algorithm was applied, and for experimental setups with the number of peptides > 50, rcBBA was used.

We next tested *copepodTCR*'s ability to achieve balanced peptide pools. We assessed balance by comparing the theoretical balance expected given the total number of peptides, pools, and peptide occurrences across pools, with the actual balance generated by the CPP scheme. *copepodTCR* consistently achieves a near-perfect balance in the distribution of peptides across pools irrespective of the total number of peptides being tested (Supplementary Fig 1C). For instance, in the above example with 16 pools, and 4 pools per peptide, the number of peptides per pool deviates by no more than 4 peptides from the optimal balance. On average, we achieve as low as 0.5 and 0.875 peptides deviation from the theoretical number of peptides per pool for pooling schemes of 50 and 950 peptides, respectively.

### The CPP scheme in *copepodTCR* enables detection of erroneously non-activated pools and narrows down the list of possible peptides.

The CPP schemes produced by *copepodTCR* ensure a constant number of activated pools for any given peptide. This feature enables the detection of experimental errors (False Positives and False Negatives) and allows narrowing down the list of possible peptides based on pools that were correctly activated (Supplementary Fig 2).

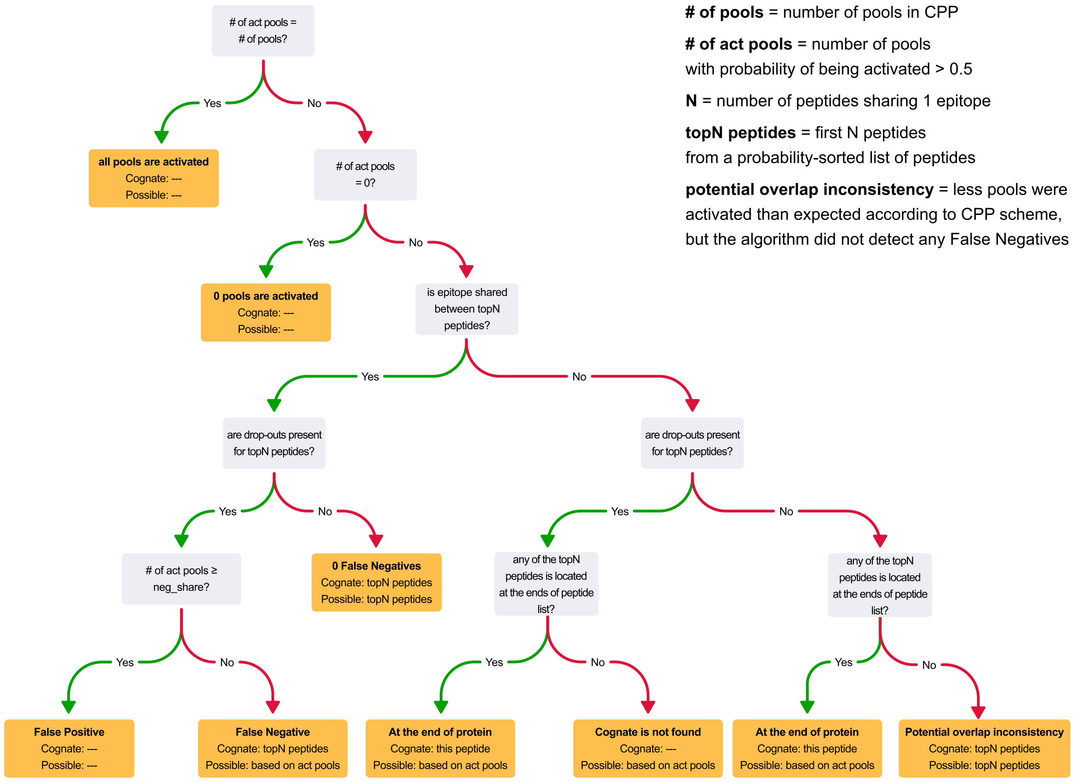


**Supplementary Figure 2. Decision diagram for identifying cognate peptides using probabilities estimated by the Bayesian model.** Using the CPP scheme, the algorithm determines pools expected to be activated for each peptide, ranks peptides based on their probability derived from the Bayesian model, and then identifies potential False Positives and False Negatives. Checks for cognate peptides located at the end of the protein are also included.

### References

1. He, G, Kovaleva, VA, Barton, C, Thomas, PG, Pogorelyy, MV, Meyer, HV & Huang, Q. Unbiased and Error-Detecting Combinatorial Pooling Experiments with Balanced Constant-Weight Gray Codes for Consecutive Positives Detection. doi:10.48550/arXiv.2502.08214 (2025).
